# Performance characteristics of next-generation sequencing for antimicrobial resistance gene detection in genomes and metagenomes

**DOI:** 10.1101/2021.06.25.449921

**Authors:** Ashley M. Rooney, Amogelang R. Raphenya, Roberto G. Melano, Christine Seah, Noelle R. Yee, Derek R. MacFadden, Andrew G. McArthur, Pierre H.H. Schneeberger, Bryan Coburn

**Author notes:** **Address correspondence to:** Bryan Coburn, University Health Network, Division of Advanced Diagnostics, Princess Margaret Cancer Research Tower, 101 College St., Toronto, ON, M5G 1L7. **Alternate corresponding author:** Pierre Schneeberger, Swiss Tropical and Public Health Institute, Department Medical Parasitology and Infection Biology, University of Basel, Socinstrasse 57. 4051 Basel, Petersplatz 1. 4001 Basel, Switzerland.

## Abstract

Short-read sequencing provides a culture-independent method for the detection of antimicrobial resistance (AMR) genes from single bacterial genomes and metagenomic samples. However, the performance characteristics of these approaches have not been systematically characterized. We compared assembly- and read-based approaches to determine sensitivity, positive predictive value, and sequencing limits of detection required for AMR gene detection in an *Escherichia coli* ST38 isolate spiked into a synthetic microbial community at varying abundances. Using an assembly-based method the limit of detection was 15X genome coverage. We are confident in AMR gene detection at target relative abundances of 100% to 1%, where a target abundance of 1% would require assembly of approximately 30 million reads to achieve 15X target coverage. Recent studies assessing AMR gene content in metagenomic samples may be inadequately sequenced to achieve high sensitivity. Our study informs future sequencing projects and analytical strategies for genomic and metagenomic AMR gene detection.

## Introduction

Increasing throughput and decreasing costs of DNA sequencing have made whole genome and metagenomic sequencing accessible for antimicrobial resistance (AMR) gene detection on a broad scale. This technology is a useful epidemiological tool^1,2^ and there are increased efforts to correlate isolate genotype with phenotypic resistance^3,4^. The ‘resistome’^5^ is the total genetic content of the microbiome with the potential to confer resistance to antibiotics, and there has been significant interest in characterizing the AMR gene content in the environment^6–8^, humans^9,10^, and other mammals^11,12^. Large trials investigating antibiotic efficacy have also included the development of AMR in the gut microbiome as an outcome^13^.

Novel methods for AMR gene detection have been developed to tackle the challenge of AMR gene identification from single isolates and metagenomic samples, generally using either assembly-based or read-based approaches, but there is currently no universal standard^14^. Notably, there are no recommendations for optimal sequencing depths required to identify AMR genes in complex metagenomic samples, and the performance characteristics of different sequencing depths using common AMR detection tools for genomes and metagenomes have not been established.

In this study, we used the Resistance Gene Identifier (RGI) and the Comprehensive Antibiotic Resistance Gene Database (CARD) and compared an assembly- and read-based approach to assess the limits of detection, sensitivity, and PPV of sequencing to detect known AMR genes in a multidrug-resistant *E. coli* isolate that represented varying abundances in a complex metagenome. We highlight the importance of maintaining minimum target genome coverage to detect AMR genes when the target organism is at varying relative abundances in a metagenomic sample and provide an estimate of minimum required sequencing depths of target organisms to maintain adequate sensitivity.

## Results

### Assembly-based AMR gene detection in *Escherichia coli* ST38

To ensure that sequencing effort was not a limiting factor in AMR gene detection, we subjected an *Escherichia coli* (*E. coli*) ST38 isolate to deep sequencing and obtained approximately 136 million reads (̴6,800X genome coverage). To simulate sequencing at lower depths, we randomly subsampled 5,000,000 (̴250X), 1,000,000 (̴50X), 500,000 (~25X), 300,000 (̴15X), 250,000 (̴12.5X), 200,000 (̴10X), 150,000 (̴7.5X), 100,000 (̴5X), 50,000 (̴2.5X), and 10,000 (̴0.5X) read pairs, and bootstrapped each subsample 100 times, with replacement, to provide confidence in the AMR genes detected in each subsample. We considered AMR genes detected with ≥90% detection frequency as high confidence genes, whereas those detected with ≤50% detection frequency were considered low confidence genes.

Using the SPAdes genome assembler, a sequencing depth of 300,000 reads or approximately 15X coverage was sufficient to detect *bla*_CTX-M-15_, and *parC* and *gyrA* single nucleotide polymorphisms (SNPs) as well as 69 other genes with greater than ≥90% detection frequency (Fig. 1a). Other resistance genes included 3 different beta-lactamases (*bla*_*TEM-1*_, *bla*_*OXA-1*_, and *bla*_*AmpC*_), 5 unique aminoglycoside transferases, and 46 distinct efflux pump genes (Fig. 1b). A lower sequencing depth was adequate to detect the SNPs with ≥90% detection frequency (150,000 reads for *gyrA* and 100,000 for *parC*) compared to *bla*_CTX-M-15_ (200,000 reads) (Fig. 1a). There were AMR genes detected less frequently (≤50% detection frequency) across all sequencing depths except at 500,000 reads where no AMR genes were detected at ≤50% detection frequency (Fig. 1a).

**Figure 1.**
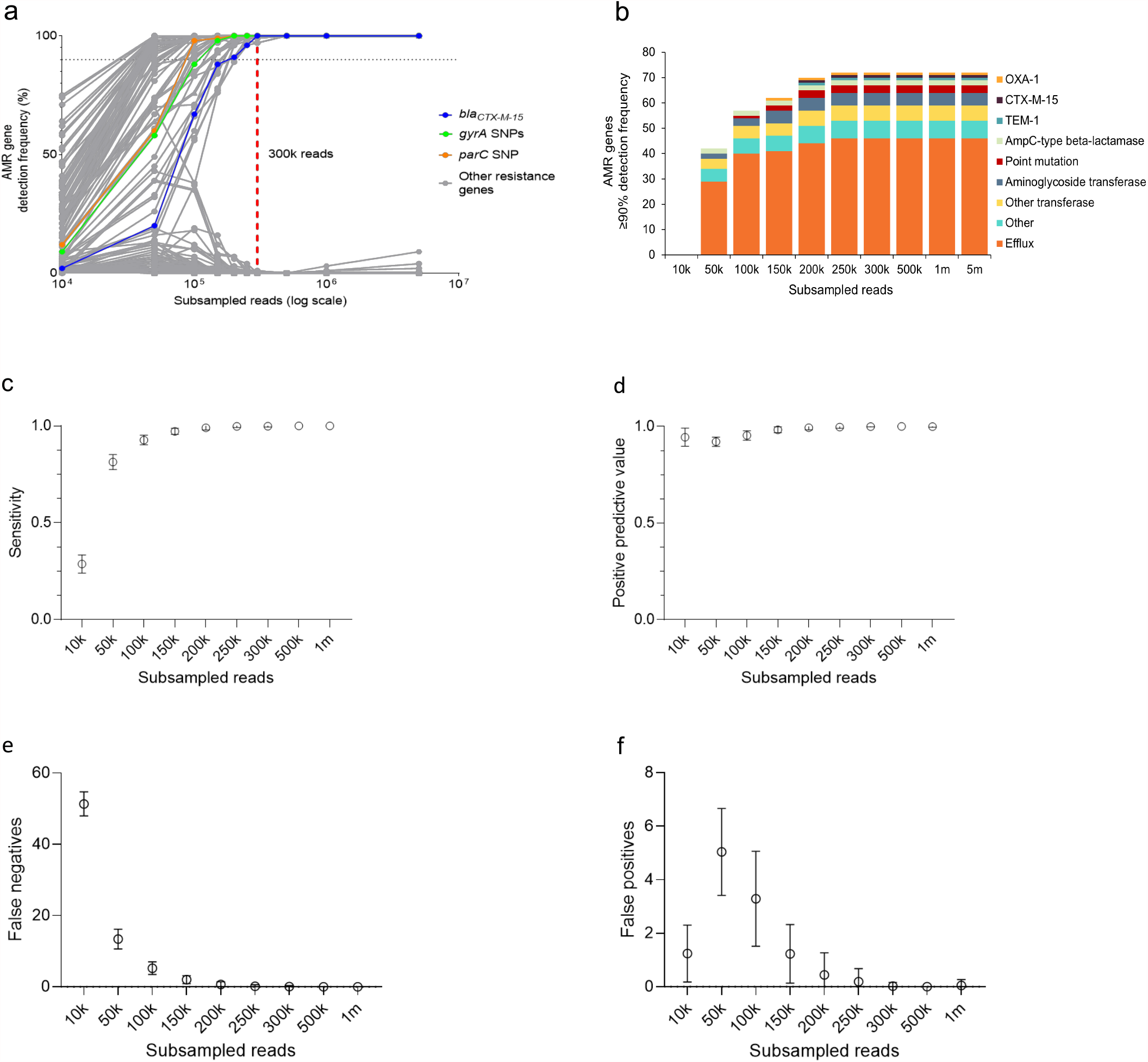
**a**, A rarefaction plot of *Escherichia coli* ST38 AMR genes detected across subsamples using an assembly-based approach. Individual dots represent a single AMR gene and are connected by lines to demonstrate trends in detection across subsamples. *bla*_*CTX-M-15*_, *gyrA* SNPs (S83L, D87N), and *parC* SNP (S80I) are highlighted as previously identified resistance determinants for this strain. The horizontal dotted line marks 90% detection frequency. The red vertical dashed line marks the subsample at 300,000 reads. **b**, Histogram of the number of unique AMR genes with ≥90% detection frequency summarized by categories detected across subsamples. (**c-f**) Performance of AMR gene classification across subsamples. A 5 million read subsample was used as reference to calculate sensitivity (**c**), positive predictive value (**d**), false negatives (**e**), and false positives (**f). c-f**, the mean and standard deviation are plotted.

To demonstrate how sequencing depth affects the performance of AMR gene detection, we used a single 5 million read subsample (̴250X coverage) as a reference to calculate sensitivity and positive predictive value (PPV) across subsamples. We did not use specificity as a metric to assess performance due to the high number of true negatives which would inflate specificity. A depth of 300,000 reads performed similarly to 1 million reads for sensitivity (1.00 ± 0.00 vs 1.00 ± 0.00, Fig. 1c) and PPV (mean = 1.00 ± 0.00 vs 1.00 ± 0.00, Fig. 1d) with low false negatives (0.09 ± 0.29, Fig. 1e) and false positives (0.02 ± 0.14, Fig. 1f) (mean and standard deviation).

The Basic Local Alignment Tool (BLAST) is a highly sensitive alignment tool in common use^15^. We aimed to compare high and low confidence AMR genes predicted using BLAST or DIAMOND, a faster alternative sequence alignment tool to BLAST^15^. Overall, BLAST predicted more AMR genes across all subsamples (Supplementary Fig. 1c). Using BLAST, as sequencing depth increased, 72 AMR genes achieved ≥90% detection frequency by 300,000 reads (Supplementary Fig. 1a), which is consistent with results using DIAMOND (Supplementary Fig. 1b). Between BLAST and DIAMOND, the genes predicted with ≥90% detection frequency at subsamples ≥300,000 reads were similar in number (approximately 72 genes were detected by both methods) as well as annotation (Supplementary Fig. 1d & 1f). Across all subsamples, more genes were predicted with ≤50% detection frequency using BLAST. For example, in the 300,000 read subsample, 2 genes were detected with ≤50% detection frequency using DIAMOND and 16 genes were predicted using BLAST (Supplementary Fig. 1e). Of the total AMR genes detected by BLAST and DIAMOND with ≤50% detection frequency, at subsamples 300,000, 500,000, 1,000,000 and 5,000,000 reads, 14/16 (87%), 4/4 (100.0%), 8/11 (73%), and 22/25 (88%), respectively, were unique to BLAST (Supplementary Fig. 1f). We proceeded to use DIAMOND for the remainder of the study as fewer low confidence genes were predicted.

### Read-based AMR gene detection in *E. coli* ST38

We next compared AMR genes predicted using a read-based approach to AMR genes predicted with an assembly-based approach in the *E. coli* ST38 isolate. We chose to use KMA over other read alignment tools as it produces a consensus sequence which can be used for SNP detection^16^. For these analyses, we compared the AMR genes predicted across subsamples using KMA to those predicted using SPAdes assemblies in the 5 million read subsample.

We found that 200,000 reads (̴10X coverage) were sufficient to identify most AMR genes using KMA read alignment with ≥90% detection frequency (Fig. 2a) with a sensitivity of 93% ± 0.0% and 5.0 ± 0.2 false negatives (mean ± standard deviation) (Fig 2c & 2e). However, there were genes that had ≤50% detection frequency at 200,000 reads including the known *gyrA* and *parC* SNPs as well as *KpnE, KpnF, bla*_*TEM-1*_, and the gene annotated as homologous to *Haemophilus* penicillin-binding protein 3 conferring resistance to beta-lactams. The AMR gene *rsmA* required 5 million reads to achieve a detection frequency of ≥90% (Fig. 2a). At 200,000 reads there were 30 additional AMR genes predicted at ≥90% detection frequency that were not detected in the reference (Fig. 2b), with a PPV of 66% ± 0.0% and 34.8 ± 1.1 false positives at this depth (Fig. 2d & 2f).

**Figure 2.**
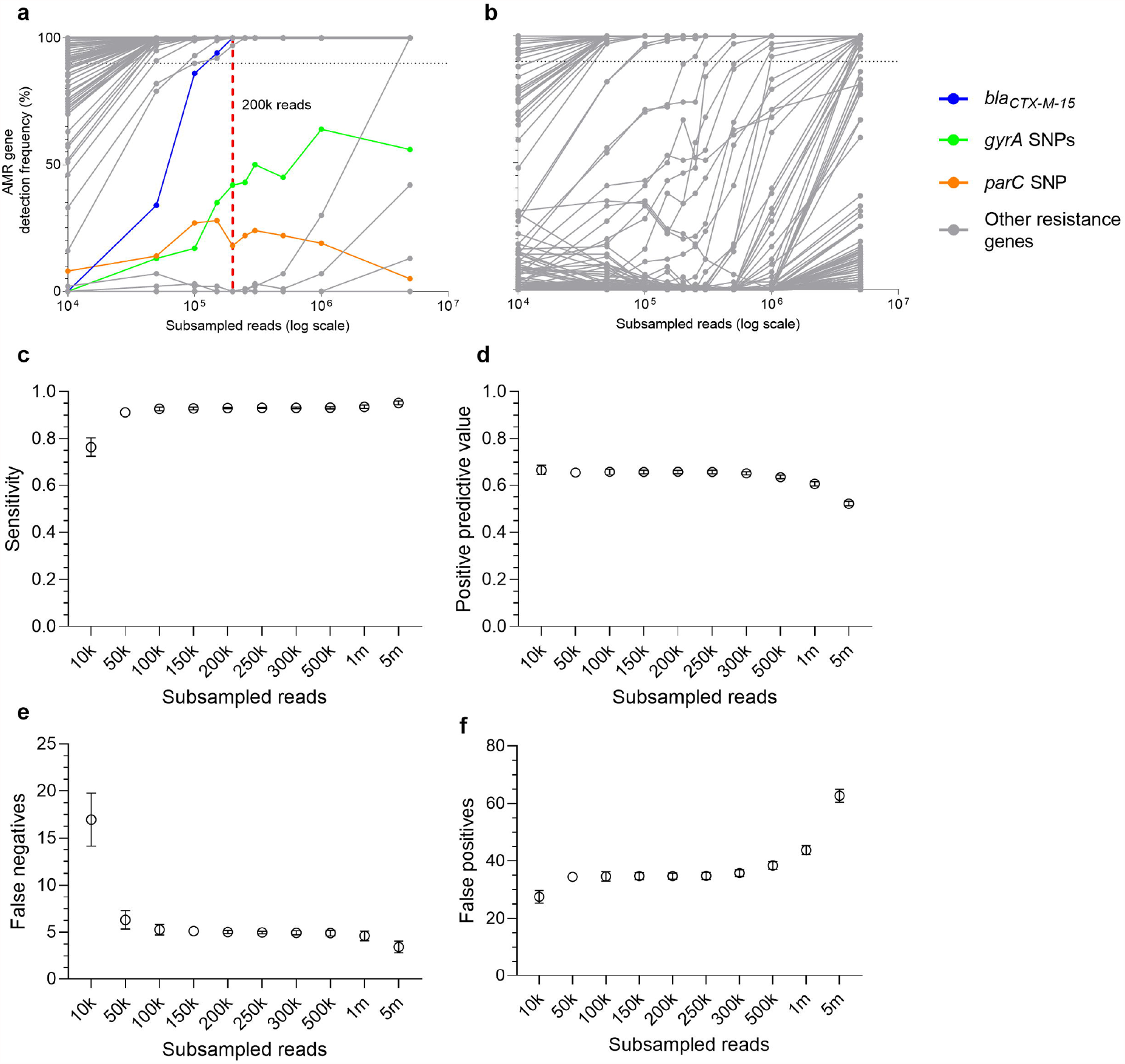
**a-b**, Rarefaction plots of *Escherichia coli* ST38 AMR genes detected across subsamples using a read-based approach. Individual dots represent a single AMR gene and are connected by lines to demonstrate trends in detection across subsamples. *bla*_*CTX-M-15*_, *gyrA* SNPs (S83L, D87N), and *parC* SNP (S80I) are highlighted as previously identified resistance determinants for this strain. The horizontal dotted line marks 90% detection frequency. The red vertical dashed line marks the subsample at 200,000 reads. **a**, AMR genes detected across subsamples which are present in the reference, **b**, AMR genes detected across subsamples which are not present in the reference. (**c-f**) Performance of AMR gene classification across subsamples, where performance was measured by sensitivity(**c**), positive predictive value (**d**), false negatives (**e**), and false positives (**f). c-f**, the mean and standard deviation are plotted. **a-e**, the reference used was all AMR genes detected using an assembly-based approach in the *E. coli* ST 38 isolate subsampled at 5 million reads.

When a read-based approach is used to identify AMR genes, strategies are often applied to improve precision and remove false positives. However, there is a trade-off between increased precision and decreased recall which is important to quantify. We assessed the effects of four AMR gene filtering strategies, at a range of cut-off values, on the performance of the 200,000 read depth for the detection of AMR genes in *E. coli* ST38 using KMA. The AMR gene filtering strategies included percent coverage, average depth of coverage, number of completely mapped reads, and the average mapping quality score (MAPQ score). The unfiltered precision and recall was 66% and 93%, respectively. No filtering strategy that we assessed significantly improved the precision, based on the precision-recall curve (precision = PPV; recall = sensitivity, Fig. 3). Out of the four strategies, percent coverage achieved the greatest increase in precision at the highest stringency cut-off of 100% allele coverage (unfiltered precision: 66%; filtered precision: 76%). However, at this cut-off, the recall decreased from 93% to 79%. When filtering by depth of coverage, completely mapped reads, or MAPQ score, the highest increase in precision was from 66% to 74% (recall: 76%), 74% (recall: 54%), and 69% (recall: 87%), respectively. As expected, as the cut-offs became more stringent for each filtering strategy, the recall decreased. Filtering based on percent coverage had the least affect on recall, even at the highest stringency cut-off (100% allele coverage) compared to the other strategies.

**Figure 3.**
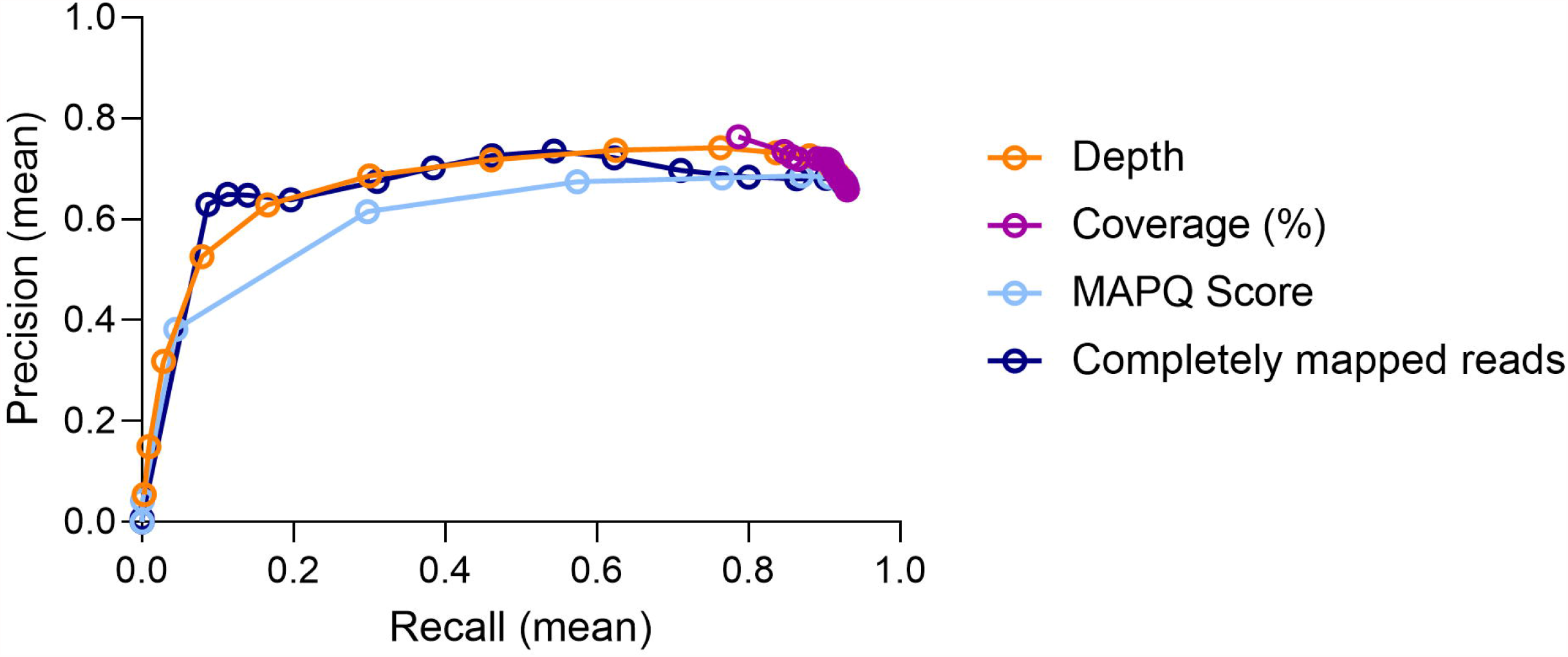
Precision-recall curve for an assessment of the effects of filtering strategies on the read-based (KMA) classification of all AMR genes in *Escherichia coli* ST38 at 200,000 reads. Precision is a measure of positive predictive value and recall is a measure of sensitivity.

### Detection of *E. coli* ST38 AMR genes at a range of relative abundances in a complex metagenomic sample

To demonstrate the effect of target organism relative abundance on AMR gene detection in a multi-species metagenome consisting of 34 bacterial species, the DNA of *E. coli* ST38 and the complex community were combined to create synthetic metagenomic samples where *E. coli* ST38 represented approximately 90%, 50%, 10%, and 1% of the total metagenome. Based on the sequencing limit of detection of 300,000 reads in the single *E. coli* ST38 isolate (100% relative abundance), we estimated that at 90%, 50%, 10%, and 1% relative abundance, 333,333, 600,000, 3,000,000 and 30,000,000 reads, respectively, would be required to detect the known AMR genes (*bla*_CTX-M-15_, *parC, gyrA* SNPs) contributed by the *E. coli* ST38 isolate with ≥90% detection frequency.

Using metaSPAdes as the metagenomic assembly tool, we observed that as the *E. coli* ST38 relative abundance decreased, the number of reads necessary to detect the *bla*_CTX-M-15_ (Fig. 4a), the *gyrA* SNPs (Fig. 4b), and the *parC* SNP (Fig. 4c), as well as the 5 aminoglycoside transferases, *bla*_*TEM-1*_, and *bla*_*OXA-1*_, increased (Supplementary Figure 2a-f). The detection rate approximated our expectations at relative abundances >1% (Fig. 4a-c). For the combined sample containing *E. coli* ST38 at 1% relative abundance, we did not detect the *gyrA* SNPs in any of the 10 bootstraps at 30,000,000 reads (Fig. 4b), while the *bla*_CTX-M-15_ and the *parC* SNP had a detection frequency of 90% (9/10 bootstraps) and 100% (10/10 bootstraps), respectively (Fig. 4a & 4c).

**Figure 4.**
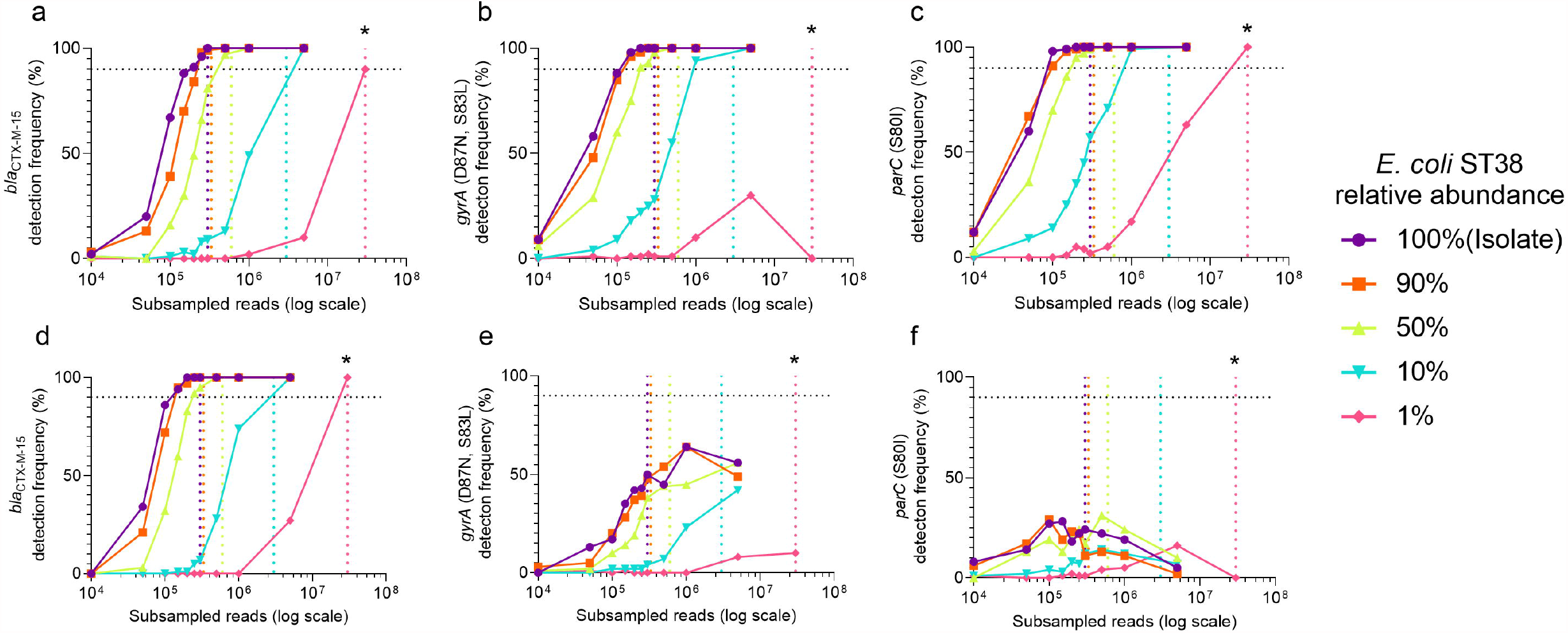
Detection of *bla*_CTX-M-15_ (**a**,**d**), *gyrA* (S83L, D87N) (**b**,**e**), and *parC* (S80I) (**c**,**f**) from *E. coli* ST38 isolate (100%) across subsamples at varying strain relative abundances (90%, 50%, 10%, 1%) in a complex (multi-species) metagenomic sample using either an assembly-based approach (metaSPAdes)(**a-c**) or a read-based approach (KMA)(**d-f**). The horizontal dotted line marks 90% detection frequency. The vertical dotted lines represent the read depths where each gene was estimated to be detected with ≥90% detection frequency for each colour matched relative abundance using an assembly-based approach. The estimated read depths are 300,000, 333,333, 600,000, 3,000,000, and 30,000,000 to detect *bla*_CTX-M-15_, *gyrA*, and *parC* SNPs in ≥90% detection frequency when *E. coli* ST38 is at 100%, 90%, 50%, 10%, and 1% relative abundance respectively. For all subsamples 100 bootstraps were performed, except for the 30,000,000 read subsample where 10 bootstraps were performed, which is marked by an asterisk (*). The 30,000,000 read subsample was performed on one combined sample where *E. coli* represented 1% relative abundance.

Although read-based approaches are often thought of as highly sensitive tools for AMR gene detection in low abundance organisms^14^, KMA did not significantly improve the limit of detection for *bla*_CTX-M-15_ (Fig. 4d), the 5 aminoglycoside transferases or *bla*_*OXA-1*_ (Supplementary Figure 3a-e & 3g) compared to assembly. The detection frequency was low for the *gyrA* and *parC* SNPs (Fig. 4e & 4f), as well as *bla*_*TEM-1*_ (Supplementary Figure 3f) across subsamples and metagenomic samples. Instead of *bla*_*TEM-1*_, KMA aligned reads to three TEM-variants with ≥90% detection frequency in at least one metagenomic sample, of which *bla*_*TEM-181*_ had the most reads aligning to the allele (Supplementary Figure 4d-f). In comparison, all TEM-variants predicted using KMA had low detection frequency using metaSPAdes (Supplementary Figure 4a-c). Sanger sequencing confirmed the presence of *bla*_*TEM-1*_ and not *bla*_*TEM-181*_ in the *E. coli* ST38 isolate. Lastly, the total number of high confidence AMR genes in the metagenomic samples was significantly higher using KMA compared to those genes detected using metaSPAdes.

### Validation of 15X coverage across *E. coli* isolates

To validate the 300,000 read depth/15X coverage threshold, we applied our assembly-based approach to 948 *E. coli* isolates^17^. The isolates were previously sequenced to an average of 100X coverage using 150-bp paired-end Illumina sequencing. We performed a subsample from each isolate at 300,000 reads and compared the AMR genes predicted at 300,000 reads to the AMR genes predicted at the original sequencing depth to calculate sensitivity, PPV, and F1 score for each isolate. The F1 score is a harmonic measure of sensitivity and PPV, where a score of 1 would indicate perfect sensitivity and PPV.

Across the *E. coli* isolate set, we observed a total of 322 unique AMR genes. Performance of the 300,000 read depth is summarized in Figure 5. The F1 score was 1 for 658/948 (69.4%) isolates. There were 290/948 (30.6%) isolates with a F1 score of <1, where 228/290 (78.6%) isolates had an F1 score between 0.99 – 0.98, 49/290 (16.9%) had an F1 score between 0.95 – 0.97, 11/290 (3.8%) had an F1 score between 0.90 – 0.94 and the remaining isolates had an F1 score 0.89 and 0.65. Of the 290 isolates with F1 score <1 (less than perfect agreement), 84 (29.0%) had a PPV of <1 and 261 (90.0%) had a sensitivity of <1. For the isolates with a PPV of <1, the median number of false positives was 1 (range 1-70). The isolate with 70 false positives is an outlier. For the isolates with a sensitivity of <1, the median number of false negatives was 1 (range 1-15).

**Figure 5.**
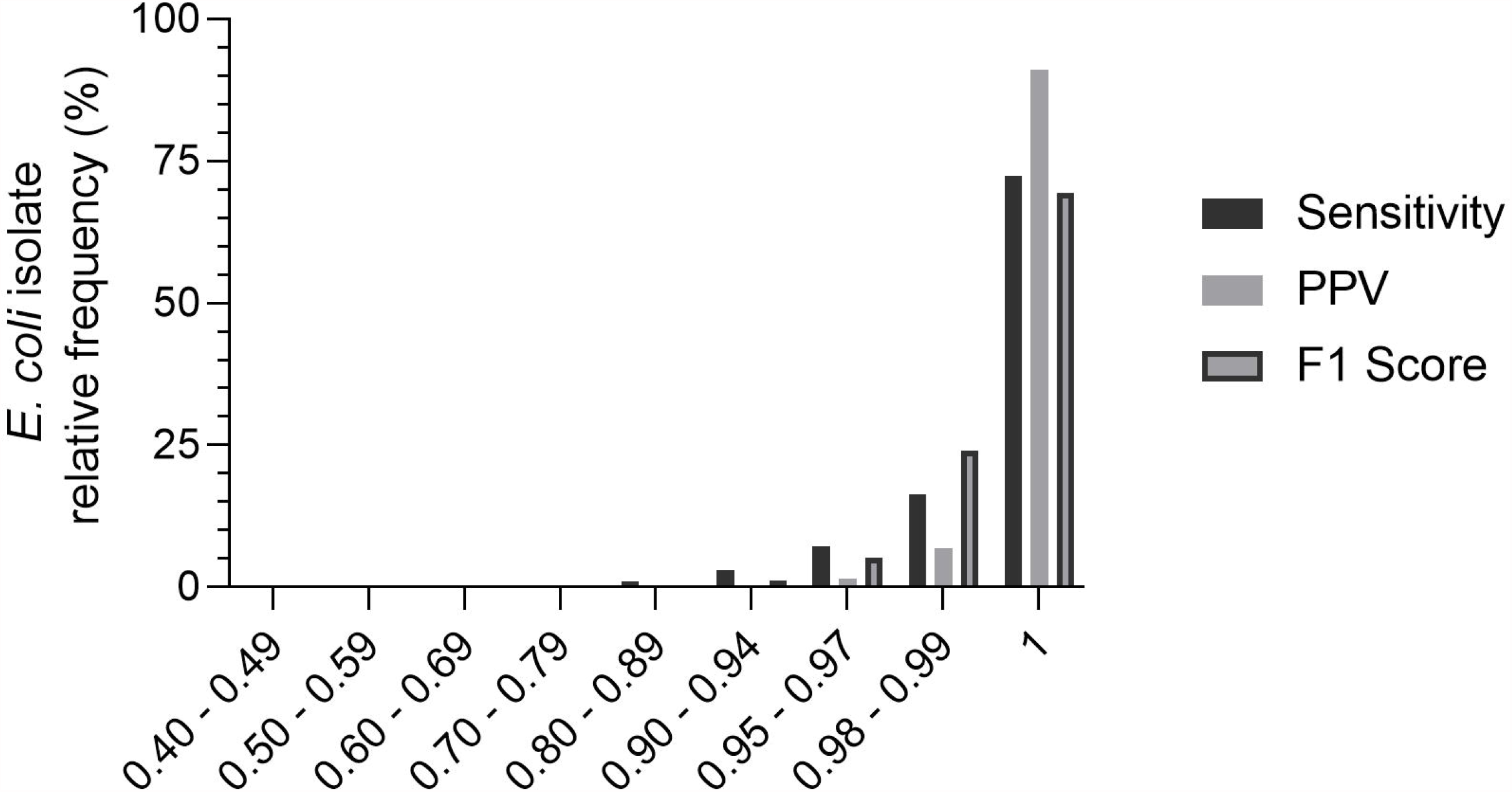
Relative frequency distribution of *E. coli* isolates (n = 948) by performance of 300,000 reads for AMR gene detection using an assembly-based approach. Performance is measured by sensitivity, positive predictive value (PPV), and F1 score.

Of the 322 unique AMR genes, 21 genes (6.5%) were classified as true positive for all 948 isolates, where 90.5% (19/21) were efflux-associated genes. The top three AMR genes that contributed the most false negatives were *APH(6)-ld* (n = 25), *sul2* (n = 23), and *mphA* (n = 23), while the top three genes that contributed the most false positives were *bla*_*OXA-320*_ (n = 13), aadA (n = 6), and *bla*_*OXA-140*_ (n = 6).

### Validation of 15X coverage in metagenomic samples

From a public dataset of 10 rectal surveillance swabs which were vancomycin resistant *Enterococcus* (VRE) positive by culture and *vanA* positive in 9/10 swabs by Illumina sequencing ^18^, we validated 15X *Enterococcus* genome coverage for the detection of *vanA*. The study authors performed 2 × 75 bp sequencing and achieved a mean 9.1 million reads (range: 5.7 million – 15 million reads), post-quality filtering and removal of human reads. The rectal swab samples had a range of *Enterococcus* relative abundances (median: 0.10; range: 0.80 – 0.0002) and genome coverages (median: 21X; range: 375X – 0.07X) (Fig. 6a). Rectal swab number 8 had the highest *Enterococcus* relative abundance of 0.80 and, due to the large number of sequencing reads (125 million), had the largest estimated target genome coverage of ̴375X. Rectal swab number 4 had the lowest *Enterococcus* relative abundance (0.0002) and 91 million sequences, which resulted in a target genome coverage of ̴0.07X for this sample.

**Figure 6.**
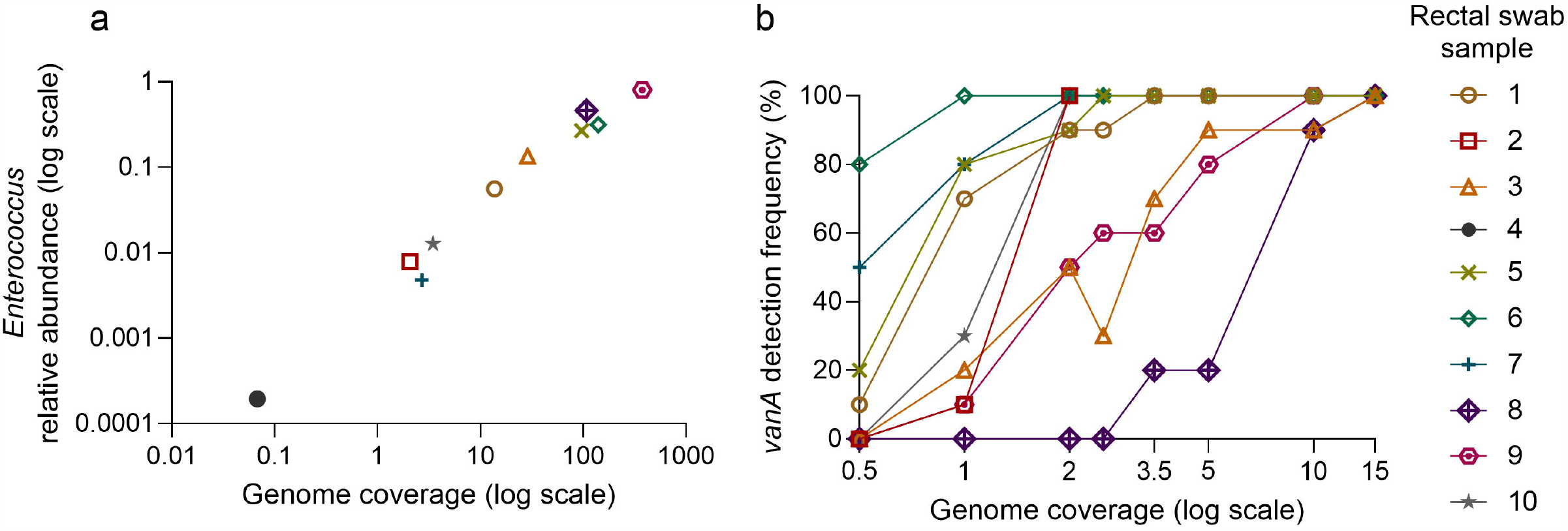
Detection of *vanA* in rectal swab samples postive for vancomycin-resistant *Enterococcus* from a public dataset. **a** *Enterococcus* relative abundance by estimated total genome coverage, each rectal swab sample is represented by an icon. **b** *vanA* detection frequency across genome coverages for each rectal swab sample. Rectal swab sample 4 is not plotted as *vanA* was not detected with the total number of sequences available. For each sample, 10 bootstraps were performed at each genome coverage depth. To calculate *Enterococcus* genome coverage, we assumed a genome size of 4 Mbp.

To assess whether a minimum of 15X target genome coverage is sufficient to detect *vanA* in the rectal swab metagenomes, we subsampled reads to achieve a range of target genome coverages from 0.5X - 15X and then bootstrapped 10 times at each subsample to determine *vanA* detection frequency. The results of this analysis are shown in Figure 6b. To achieve 100% detection frequency of the *vanA* gene across rectal swab samples, 5 samples required *Enterococcus* genome coverage of less than 5X (rectal swabs 1,5-7, and 10), while 2 required at least 15X coverage (rectal swabs 3 and 8). At 15X *Enterococcus* genome coverage, *vanA* was detected in 10/10 bootstraps for all samples that had adequate sequencing depth for subsampling. Rectal swab number 4 did not have enough reads to achieve 0.5X *Enterococcus* genome coverage and *vanA* was not detected when we analyzed all reads available, which is consistent with the authors’ published findings that describe their inability to detect *vanA* using paired-end Illumina sequencing^18^.

### Estimates of the sensitivity of sequencing depth for AMR gene detection in published data sets

Recent publications assessing AMR gene content in metagenomic samples may not have achieved optimal sensitivity for AMR gene detection if they were to use a contig assembly approach. As we have observed, the relative abundance of the target organism affects sensitivity to detect AMR genes in a metagenomic sample. We gathered sequencing depths and read length information from three recently published studies that reported AMR genes in metagenomic samples. Study 1 assessed and compared the resistome of 1174 gut and oral samples from previously published sources distributed by country^9^. For our analyses, we included 1132/1174 from Study 1 for which we had complete read length data (excluding 42/1174, 3.6%). Study 2 performed a longitudinal assessment of the gut microbiota and resistome of healthy veterinary students exposed to a Chinese swine farm environment. A total 63 metagenomic samples were sequenced which consisted of human stool and environmental samples^6^. Study 3 was conducted in Denmark and evaluated the changes in the gut microbiota composition and resistome of 12 healthy male volunteers before and after antimicrobial exposure^10^. A total of 57 stool samples were subject to metagenomic analyses. Studies 1 and 3 used a read-based approach for AMR gene prediction, while Study 2 used an assembly-based approach.

We were interested in estimating AMR gene detection frequency from the published sample sequencing depths for a hypothetical target organism (presumed genome size 6 Mbp) at a range of potential relative abundances. Assuming detection frequency is related to sequencing sensitivity, we calculated coverage as an estimate of sequencing depth and interpolated detection frequency values from a sigmoidal curve fit to the *E. coli* ST38 *bla*_CTX-M-15_ detection data as seen in Figure 4a.

As the relative abundance of the hypothetical target organism decreased, more sequencing effort was required to achieve 100% estimated detection frequency of all AMR genes (Fig. 7). Most published samples had achieved ≥95% estimated detection frequency for all AMR genes for a target organism at relative abundance of 100% (1251/1252; 99.9%), 90% (1250/1252; 99.8%) and 50% (1247/1252; 99.6%). However, the proportion of samples with at least 90% estimated detection frequency was lower for a target organism relative abundance of 10% (1090/1252; 87.1%) and 1% (454/1252; 36.3%). Additionally, 29.5% (369/1252) of samples were not sequenced sufficiently to achieve >50% estimated detection frequency for a target organism relative abundance of 1%, where 9.2% (115/1252) had less than 1% estimated detection frequency (Fig. 7), suggesting that in these studies, sequencing depth may be inadequate to achieve a high sensitivity for detection of AMR genes in low abundance organisms.

**Figure 7.**
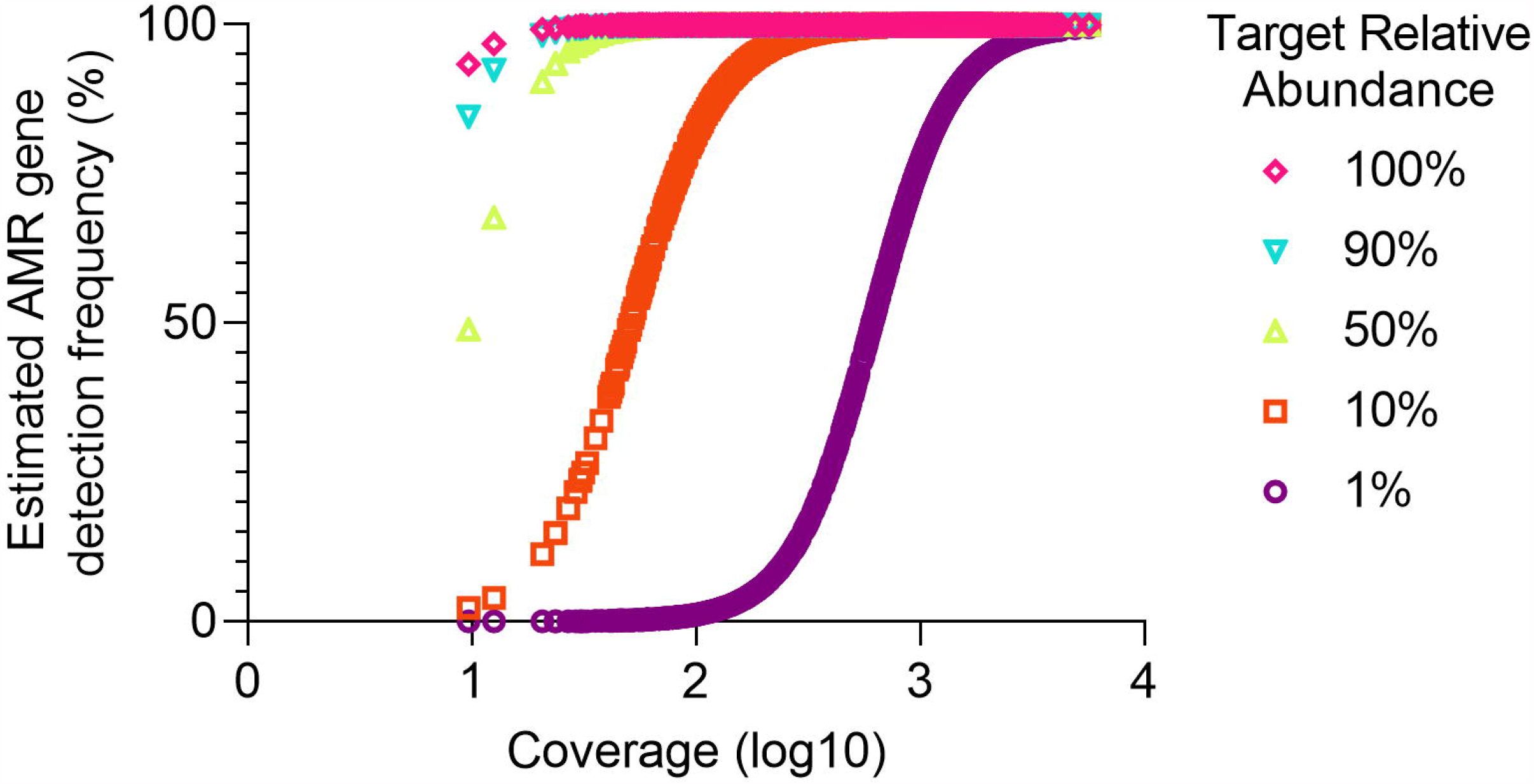
Estimated AMR gene detection frequency by sample coverage of a hypothetical target organism at different relative abundances. Sample sequencing information was extracted from three published datasets (n = 1252). Detection frequency was estimated by interpolating values from sigmoidal curves fit to the *bla*_CTX-M-15_ data from samples where *E. coli* represented 100%, 90%, 50%, 10%, and 1% relative abundance. Coverage was estimated assuming a target genome size of 6 Mbp.

## Discussion

Our goal was to characterize the performance (sensitivity, PPV, and limits of detection) of genomic and metagenomic approaches for the detection of known predictors of AMR in microbes to inform the use of these approaches in human, animal, and environmental studies. It is axiomatic that sequencing depth affects AMR gene assay sensitivity in single isolates^19,20^ and within a microbiome^21^, however, the performance characteristics of sequencing have not been systematically assessed. In published reports, a range of whole genome sequencing depths for single isolates, from 30X coverage up to 100X coverage, are often used to define quality control limits, but these are not considered standard^3,19,22,23^. Estimating the coverage of the metagenome required to ensure high sensitivity is not a new concept^24^, but we are unaware of a study that attempts to precisely quantify sequencing depths required to detect AMR genes in a target organism across varying relative abundances in mixed metagenomes using standard methods.

Using a *de novo* contig-assembly approach, we found that approximately 15X coverage (300,000, 2 × 150 bp paired-end reads of a 6 Mbp genome) provides similar sensitivity to higher sequencing depths for the detection of AMR genes in *E. coli* isolates and is sufficient for detecting SNPs and other resistance genes. Although sequencing depths as low as 0.5 million reads have been proposed to capture the total compositional information of metagenomes^25^, greater sequencing depth is required for the detection of AMR genes in organisms with low relative abundance, which can require as many as 30 million reads to achieve adequate sensitivity for organisms at a relative abundance of 1%.

For some study purposes, detection of AMR genes in low abundance organisms may be critical for study interpretation. Human observational studies have demonstrated that pathogens at both high and low relative abundances in complex gut microbial communities are associated with subsequent infections or death. Dominance of a microbial community by a pathogen is associated with subsequent infection^26–28^, but even at relative abundances as low as 1% - 0.1%, pathogens detected in stool have been implicated in subsequent bacteremia in hematopoietic stem cell transplant recipients^29^, as well as bacteriuria and urinary tract infection^30^, indicating that detection of AMR genes may be clinically significant even at very low relative abundance thresholds. Based on our findings, approximately 64% of the samples in recent studies evaluating AMR gene content in the metagenome are not sequenced at a sufficient depth to detect AMR genes in a target organism at 1% relative abundance. Thus, potentially clinically meaningful resistance determinants may not be detected with common sequencing depths such as those we analyzed in published studies.

Assembly is time-consuming, requires large amounts of computing power for metagenomic samples, and may also contribute to loss of data^31^. Alternative approaches to assembly such as read alignment^16,32–34^ and kmer-based approaches^31^ may require less sequencing information for AMR gene detection, which is useful for detecting AMR genes in low abundance organisms in complex communities. Compared to assembly, our read-based approach (KMA) did not significantly improve the limit of detection in *E. coli* ST38 or metagenomes, even for a low abundance target, and at times suffered from the AMR allele network problem^35^, where reads from a single gene were aligned to multiple closely related reference alleles, e.g. *bla*_*TEM-181*_, despite its improvements over other read alignment tools for this issue^16^. Additionally, some genes (e.g. *gyrA* and *parC* SNPs and *bla*_*TEM-1*_) that had a high detection frequency with assembly, had a low detection frequency with KMA and there were a large number of false positive genes that could not be filtered without affecting the overall sensitivity of the assay. Yet, our comparison of BLAST and DIAMOND with open reading frames predicted from assembled contigs illustrated that attention should also be paid to local alignment algorithm choice when using assembly-based approaches, as DIAMOND generated less low confidence predictions.

A large majority of the AMR genes detected in the *E. coli* ST38 isolate and across the *E. coli* isolate set were efflux-associated AMR genes. Efflux-associated genes are of uncertain relevance for prediction of microbial phenotypes^36,37^. In some scenarios (e.g. human infections) it may be advantageous to limit AMR gene detection to acquired mechanisms of resistance by using a database such as ResFinder^38^, or by filtering efflux-associated genes from the total AMR genes detected, which has been previously shown to improve predictions of isolate antimicrobial susceptibility using CARD^37^. Accurate prediction of plasmids, often associated with clinically important AMR genes, remains difficult as short-read Illumina sequencing provides highly accurate base calling, but repetitive DNA regions complicate genome assembly resulting in fragmented, short contigs^39^. AMR genes may be split between multiple contigs ^40^ leaving plasmid sequences obscured^41^. Long-read sequencing technology provides a promising alternative to short-read sequencing and can overcome the issue of fragmented contig assembly^39^.

Our approach has the following limitations. We modelled a single approach utilizing a widely used sequencing strategy, two bioinformatic pipelines and one AMR detection platform (CARD) for a single organism (*E. coli*). These selections were made to reflect dominant modes of metagenome analysis in a clinically relevant organism to define the ‘order of magnitude’ of depth required for AMR gene detection from metagenomes, which may not be generalizable to all organisms, community types or modes of resistance. A main limitation of AMR gene prediction from sequencing data is the chosen database that can potentially increase false negatives. However, CARD is widely used, updated on a monthly basis, and is representative of known AMR gene diversity, especially for well-characterized pathogens such as *E. coli*^42^. Human metagenomic samples often have human DNA that can account for a large proportion of the total sample, which impacts sequencing strategies^43,44^. An understanding of the total genetic material contributed by human reads prior to sequencing would further inform sequencing effort required to maintain a minimum sequencing depth for AMR gene detection.

We have quantified sequencing depths needed to detect AMR genes in *E. coli* whole genomes and in *E. coli* from high to low relative abundances among a complex community. A minimum of 15X coverage is needed for the detection of AMR genes in *E. coli* using our AMR gene identification approach. For metagenomic samples, 15X coverage is also sufficient to detect known AMR genes in *E. coli*, but the number of sequences must increase proportionally to the decrease in relative abundance of the target organism. We believe that this analysis provides a robust benchmarking of sequencing effort for metagenomic studies in which detection of resistance is a specified outcome.

## Methods

### Sample preparation and sequencing

From a collection of previously characterized *E. coli* isolates^17^, we selected a multidrug-resistant *E. coli* ST38 with an extended spectrum beta-lactamase (*bla*_*CTX-M-15*_) and resistance-conferring SNPs in *parC* (S80I) and *gyrA* (S83L, D87N). Briefly, *E. coli* ST38 isolate was cultured from a glycerol stock on LB agar and a single colony was inoculated into 25 ml of LB broth, which was placed on a shaker incubator (130 rpm) at 37°C for 4 hours until media was turbid. Turbid media (25 ml) was transferred to a 50 ml conical tube, subject to centrifugation at 2500 g, the supernatant removed, and the pellet re-suspended in 500 μl of LB broth. A description of the complex community (MET-1) preparation was described previously^45^. Aliquots of MET-1 were stored at -80°C prior to use.

DNA was extracted from thawed MET-1 (250 μl) and the *E. coli* isolate in LB broth (250 μl) using the DNeasy PowerSoil kit (Qiagen) and DNA concentration was measured using a Qubit Fluorometer (Thermo Fisher), following the manufacturer’s instructions, respectively. *E. coli* and MET-1 DNA were combined to a final concentration of 20.1 ng/μl, while varying the concentration of *E. coli* so that it approximately represented 90%, 50%, 10%, 1%, 0.1%, 0.01%, 0.001%, and 0.0001% relative to MET-1. Sequencing libraries were prepared using the Nextera DNA Flex kit (Illumina) following the manufacturer’s instructions and stored at -20°C. All 10 samples (the *E. coli* ST38 isolate, MET-1, and 8 combined samples) were subject to paired-end sequencing at 2 × 150 bp on the NovaSeq 6000 at the Princess Margaret Genomics Centre. Since we did not achieve the minimum sequencing depth needed to detect AMR genes in the combined samples where *E. coli* ST38 represented 0.1%, 0.01%, 0.0001%, and 0.0001% we did not analyze these samples in our study.

### Bioinformatic analyses

From each pair of fastq files, Seqtk^46^ was used to subsample *n* number of reads. At each subsample, bootstrapping was performed (sampling with replacement) 100 times unless otherwise stated, where all 100 bootstraps of a subsample had a unique seed number to ensure every bootstrap was a random sampling of reads. Paired-end fastq files (read 1 and read 2) were assessed for quality using FastQC^47^. Nextera adapters were removed with Trimmomatic^48^v.0.39. Reads for *E. coli* genomes as well as MET-1 and the combined samples were assembled into contigs using SPAdes^49^ v.3.13.1, specifying the *--careful* flag, and metaSPAdes^50^ v.3.13.1, respectively, using the recommended kmer lengths 21, 33, 55, and 77. We used Metaphlan2^51^v.2.9.21 to confirm sample taxonomy, including the identity of all *E. coli* isolates and the relative abundance of *Enterococcus* species in the validation sets, respectively.

To predict AMR genes from contigs, we used RGI *main* v.5.1.0 of the Comprehensive Antibiotic Resistance Database on default settings (perfect and strict hits identified only)^42^. We specified DIAMOND^15^ v.0.8.36, or BLAST^52^ v.2.9.0 (where stated) to perform local alignment of Prodigal-predicted genes within contigs against CARD v.3.1.0^42,53^. For metagenome assembled contigs, we specified the *–low_quality* flag in RGI *main* to allow prediction of partial open reading frames by Prodigal.

To predict AMR genes from raw reads, we used KMA^16^ v.1.3.8 within RGI *bwt* v.5.2.0 to align reads to CARD. To predict AMR-conferring SNPs in the *parC* (S80I) and *gyrA* (S83L, D87N) genes, we extracted the consensus sequences generated from these read alignments and used RGI *main* v.5.2.0.

### Quantification of AMR genes

To quantify the occurrence of the *parC* (S80I) and *gyrA* (S83L, D87N) SNPs in bootstrapped subsamples of the *E*.*coli* ST38 isolate and combined samples, we extracted the individual accession numbers (from contig results only) and SNP information (contigs and raw reads) for each gene from all RGI or KMA output files. The *parC* and *gyrA* SNPs were considered present if the mutated amino acid residues S80I in *parC* and S83L, D87N in *gyrA* were correctly predicted by RGI. If RGI predicted other mutated amino acid residues in the *parC* and *gyrA* genes or did not identify any mutated amino acid residues in the *parC* and *gyrA* gene, we did not consider these SNPs as present. For all other resistance genes, we extracted unique genes from the “Best_Hit_ARO” (contigs) or “ARO_Term” (raw reads) column of each sample RGI output file, to create a new “unique AMR genes” file for each sample. Then, using Metaphlan2 v.2.9.14 we used the merge_metaphlan_tables.py command to merge the “unique AMR genes” files together, where the first column outlined the AMR genes predicted for all samples and the first row indicated the sample names. AMR gene presence was indicated by 1 and absence indicated by 0. Merging the RGI output files allowed us to quantify the frequency at which individual AMR genes were present across samples. We considered an AMR gene present in a bootstrap if the gene occurred at least once. The number of AMR genes present across bootstrapped subsamples was visualized with rarefaction curves and plotted using GraphPad Prism version 9.

### Coverage estimation

Sequencing coverage was estimated using the Lander-Waterman equation^54^. For *E. coli* ST38, we assumed a genome size of 6 Mbp. To estimate the number of reads required to detect *E. coli* ST38 at a range of relative abundances, the minimum read requirement (300,000 reads) was divided by the target relative abundance. For example, if the target relative abundance was 10%, 300,000/0.10 would equal 3,000,000 reads.

### Performance analyses

The performance of AMR gene classification was calculated using sensitivity and positive predictive value. For the single *E. coli* ST38 isolate, the AMR genes predicted in each bootstrapped subsample using a contig assembly approach with SPAdes or a read-based approach with KMA were compared to the AMR genes predicted in a subsample assembled into contigs at 5 million reads (reference). If the AMR gene was present in the bootstrap and reference, this gene was considered a true positive. If an AMR gene was not present in neither the bootstrap nor the reference, this gene was considered a true negative. False positive AMR genes were present in the bootstrap but absent in the reference and false negative AMR genes were absent in the bootstrap but present in the reference. For each bootstrap sample, the true positives, true negatives, false positives, and false negatives were summed and sensitivity and PPV were calculated.

### Validation from external datasets

To validate the performance of a 300,000 read depth across a set of *E. coli* isolates^17^, we subsampled 300,000 reads, once, from each isolate, assessed quality with FastQC and discarded isolates that failed per base sequence quality. We then compared the AMR genes detected at 300,000 reads to the AMR genes detected from the original sequence depth. We summed the true positives, true negatives, false positives, and false negatives for each isolate, then calculated sensitivity, PPV and F1 score as a balanced measure of sensitivity and PPV.

To validate 15X target genome coverage in metagenomic samples^18^ and to demonstrate *vanA* detection frequency across a range of *Enterococcus* genome coverages (0.5X – 15X), we subsampled each metagenomic sample and bootstrapped each subsample 10 times. Each subsample depth was calculated using the Lander-Waterman equation, as described above, while accounting for the *Enterococcus* relative abundance in the sample, as determined using Metaphlan2. We assumed an *Enterococcus* genome size of 4 Mbp.

### AMR gene detection frequency assessment of published datasets

We extracted the sequence depths after quality processing that were provided in each studies’ supplementary material for Study 1^9^, 2^6^ and 3^10^. For Study 3, we used the sequences reported under the heading “After human contamination removal” under the sub-heading “read-pairs”. Sequencing read lengths were reported in Study 2 (2 × 150 base pairs) and 3 (2 × 100 base pairs), but for Study 1 we extracted the read lengths from the individual studies referenced within the paper. We then estimated coverage for each sample from the published datasets and for each subsample performed on the samples where *E. coli* ST38 represented 100%, 90%, 50%, 10%, and 1% relative abundance, assuming a genome length of 6 Mbp, then log-transformed these values. Using GraphPad Prism version 9.1.2, sigmoidal curves were fit to detection frequency data for the *bla*_CTX-M-15_ for each sample where *E. coli* ST38 represented 100%, 90%, 50%, 10%, and 1% relative abundance. The equations were constrained at 0 and 100 and detection frequency was interpolated for relative abundances 100%, 90%, 50%, 10%, and 1% based on coverage estimation.

## Data availability

The dataset generated during the current study are available in the NCBI sequence read archive under the accession number PRJNA649958.

## Competing interests

The authors declare that they have no competing interests.

## Author Contributions

AMR, PHHS, and BC conceived and designed the study. BC supervised the overall study. AMR with the guidance and supervision of PHHS performed sample preparation and bioinformatic analyses. ARR provided feedback on and performed some bioinformatic analyses. RM and CS provided qPCR and Sanger sequencing support. RM, DM, and AM provided feedback during analyses. NY provided help with the bioinformatic analyses. AMR generated the figures and wrote the manuscript. All authors provided feedback during manuscript preparation and have read and approved the final manuscript.

## Acknowledgments

We greatly appreciate Dr. Emma Allen-Vercoe from the University of Guelph for sending us aliquots of the complex community, MET-1. This research was funded by the Canadian Institutes of Health Research (PJT-156214 to AM) and a David Braley Chair in Bioinformatics to AM. Some computer resources were supplied by the McMaster Service Lab and Repository computing cluster, funded in part by grants to AM from the Canada Foundation for Innovation (34531) and hardware donations for Cisco Systems Canada, Inc.

## Notes

### Competing Interest Statement

The authors have declared no competing interest.

## References

1. Jackson, B. R. et al. Implementation of Nationwide Real-time Whole-genome Sequencing to Enhance Listeriosis Outbreak Detection and Investigation. Clin. Infect. Dis. 63, 380–386 (2016).

2. Harrison, L. H. et al. Outbreak of Vancomycin-resistant Enterococcus faecium in Interventional Radiology: Detection Through Whole-genome Sequencing-based Surveillance. Clin. Infect. Dis. 70, 2336–2343 (2020).

3. Macfadden, D. R. et al. Comparing Patient Risk Factor-, Sequence Type-, and Resistance Locus Identification-Based Approaches for Predicting Antibiotic Resistance in Escherichia coli Bloodstream Infections. J. Clin. Microbiol. 57, e01780–18 (2019).

4. Břinda, K. et al. Rapid inference of antibiotic resistance and susceptibility by genomic neighbour typing. Nat. Microbiol. 5, 455–464 (2020).

5. D’Costa, V. M., McGrann, K. M., Hughes, D. W. & Wright, G. D. Sampling the antibiotic resistome. Science. 311, 374–377 (2006).

6. Sun, J. et al. Environmental remodeling of human gut microbiota and antibiotic resistome in livestock farms. Nat. Commun. 11, 1427 (2020).

7. Chng, K. R. et al. Cartography of opportunistic pathogens and antibiotic resistance genes in a tertiary hospital environment. Nat. Med. 26, 941–951 (2020).

8. Pehrsson, E. C. et al. Interconnected microbiomes and resistomes in low-income human habitats. Nature 533, 212–216 (2016).

9. Carr, V. R. et al. Abundance and diversity of resistomes differ between healthy human oral cavities and gut. Nat. Commun. 11, 693 (2020).

10. Palleja, A. et al. Recovery of gut microbiota of healthy adults following antibiotic exposure. Nat. Microbiol. 3, 1255–1265 (2018).

11. Lim, S.-K., Kim, D., Moon, D.-C., Cho, Y. & Rho, M. Antibiotic resistomes discovered in the gut microbiomes of Korean swine and cattle. Gigascience 9, 1–11 (2020).

12. Liu, J. et al. The fecal resistome of dairy cattle is associated with diet during nursing. Nat. Commun. 10, 4406 (2019).

13. Doan, T. et al. Gut microbiome alteration in MORDOR I: a community-randomized trial of mass azithromycin distribution. Nat. Med. 25, 1370–1376 (2019).

14. Boolchandani, M., D’Souza, A. W. & Dantas, G. Sequencing-based methods and resources to study antimicrobial resistance. Nat. Rev. Genet. 20, 356–370 (2019).

15. Buchfink, B., Xie, C. & Huson, D. H. Fast and sensitive protein alignment using DIAMOND. Nat. Methods 12, 59–60 (2015).

16. Clausen, P. T. L. C., Aarestrup, F. M. & Lund, O. Rapid and precise alignment of raw reads against redundant databases with KMA. BMC Bioinformatics 19, 307 (2018).

17. Macfadden, D. R. et al. Using Genetic Distance from Archived Samples for the Prediction of Antibiotic Resistance in Escherichia coli. Antimicrob. Agents Chemother. 64, e02417–19 (2020).

18. Yee, R. et al. Metagenomic next-generation sequencing of rectal swabs for the surveillance of antimicrobial-resistant organisms on the Illumina Miseq and Oxford MinION platforms. Eur. J. Clin. Microbiol. Infect. Dis. 40, 95–102 (2021).

19. Doyle, R. M. et al. Discordant bioinformatic predictions of antimicrobial resistance from whole-genome sequencing data of bacterial isolates: an inter-laboratory study. Microb. Genomics 6, e000335 (2020).

20. Cooper, A. L. et al. Systematic Evaluation of Whole Genome Sequence-Based Predictions of Salmonella Serotype and Antimicrobial Resistance. Front. Microbiol. 11, 549 (2020).

21. Zaheer, R. et al. Impact of sequencing depth on the characterization of the microbiome and resistome. Sci. Rep. 8, 5890 (2018).

22. Ellington, M. J. et al. The role of whole genome sequencing in antimicrobial susceptibility testing of bacteria: report from the EUCAST Subcommittee. Clin. Microbiol. Infect. 23, 2– 22 (2017).

23. Mcdermott, P. F. et al. Whole-Genome Sequencing for Detecting Antimicrobial Resistance in Nontyphoidal Salmonella. Antimicrob. Agents Chemother. 60, 5515–5520 (2016).

24. Rodriguez-R, L. M. & Konstantinidis, K. T. Estimating coverage in metagenomic data sets and why it matters. ISME J. 8, 2349–2351 (2014).

25. Hillmann, B. et al. Evaluating the Information Content of Shallow Shotgun Metagenomics. mSystems 3, e00069–18 (2018).

26. Taur, Y. et al. Intestinal domination and the risk of bacteremia in patients undergoing allogeneic hematopoietic stem cell transplantation. Clin. Infect. Dis. 55, 905–914 (2012).

27. Stewart, C. J. et al. Longitudinal development of the gut microbiome and metabolome in preterm neonates with late onset sepsis and healthy controls. Microbiome 5, 75 (2017).

28. Zhai, B. et al. High-resolution mycobiota analysis reveals dynamic intestinal translocation preceding invasive candidiasis. Nat. Med. 26, 59–64 (2020).

29. Tamburini, F. B. et al. Precision identification of diverse bloodstream pathogens in the gut microbiome. Nat. Med. 24, 1809–1814 (2018).

30. Magruder, M. et al. Gut uropathogen abundance is a risk factor for development of bacteriuria and urinary tract infection. Nat. Commun. 10, 5521 (2019).

31. Clausen, P. T. L. C., Zankari, E., Aarestrup, F. M. & Lund, O. Benchmarking of methods for identification of antimicrobial resistance genes in bacterial whole genome data. J. Antimicrob. Chemother. 71, 2484–2488 (2016).

32. Langmead, B. & Salzberg, S. L. Fast gapped-read alignment with Bowtie 2. Nat. Methods 9, 357–359 (2012).

33. Li, H. & Durbin, R. Fast and accurate short read alignment with Burrows–Wheeler transform. Bioinformatics 25, 1754–1760 (2009).

34. Inouye, M. et al. SRST2: Rapid genomic surveillance for public health and hospital microbiology labs. Genome Med. 6, 90 (2014).

35. Lanza, V. F. et al. In-depth resistome analysis by targeted metagenomics. Microbiome 6, 11 (2018).

36. Moran, R. A., Anantham, S., Holt, K. E. & Hall, R. M. Prediction of antibiotic resistance from antibiotic resistance genes detected in antibiotic-resistant commensal Escherichia coli using PCR or WGS. J. Antimicrob. Chemother. 72, 700–704 (2017).

37. Mahfouz, N., Ferreira, I., Beisken, S., von Haeseler, A. & Posch, A. E. Large-scale assessment of antimicrobial resistance marker databases for genetic phenotype prediction: A systematic review. J. Antimicrob. Chemother. 75, 3099–3108 (2020).

38. Zankari, E. et al. Identification of acquired antimicrobial resistance genes. J. Antimicrob. Chemother. 67, 2640–2644 (2012).

39. Goldstein, S., Beka, L., Graf, J. & Klassen, J. L. Evaluation of strategies for the assembly of diverse bacterial genomes using MinION long-read sequencing. BMC Genomics 20, 23 (2019).

40. Su, M., Satola, S. W. & Read, T. D. Genome-Based Prediction of Bacterial Antibiotic Resistance. J. Clin. Microbiol. 57, e01405–18 (2019).

41. Robertson, J. & Nash, J. H. E. MOB-suite: software tools for clustering, reconstruction and typing of plasmids from draft assemblies. Microb. Genomics 4 (2018). doi:10.6084/m9.figshare.6177188

42. Alcock, B. P. et al. CARD 2020: antibiotic resistome surveillance with the comprehensive antibiotic resistance database. Nucleic Acids Res. 48, D517–D525 (2019).

43. Pereira-Marques, J. et al. Impact of Host DNA and Sequencing Depth on the Taxonomic Resolution of Whole Metagenome Sequencing for Microbiome Analysis. Front. Microbiol. 10, 1277 (2019).

44. Cho, M. Y. et al. Two-target quantitative PCR to predict library composition for shallow shotgun sequencing. bioRxiv 1–14 (2020). doi:10.1101/2020.09.21.304006

45. Petrof, E. O. et al. Stool substitute transplant therapy for the eradication of Clostridium difficile infection: ‘RePOOPulating’ the gut. Microbiome 1, 3 (2013).

46. Li, H. Seqtk: Toolkit for processing sequences in FASTA/Q formats. Github (2012). Available at: https://github.com/lh3/seqtk.

47. Andrews, S. FastQC: a quality control tool for high throughput sequence data. (2010).

48. Bolger, A. M., Lohse, M. & Usadel, B. Trimmomatic: a flexible trimmer for Illumina sequence data. Bioinformatics 30, 2114–2120 (2014).

49. Bankevich, A. et al. SPAdes: A New Genome Assembly Algorithm and Its Applications to Single-Cell Sequencing. J. Comput. Biol. 19, 455–477 (2012).

50. Nurk, S., Meleshko, D., Korobeynikov, A. & Pevzner, P. A. MetaSPAdes: A new versatile metagenomic assembler. Genome Res. 27, 824–834 (2017).

51. Truong, D. T. et al. MetaPhlAn2 for enhanced metagenomic taxonomic profiling. Nat. Methods 12, 902–903 (2015).

52. Camacho, C. et al. BLAST+: Architecture and applications. BMC Bioinformatics 10, 421 (2009).

53. Hyatt, D. et al. Prodigal: Prokaryotic gene recognition and translation initiation site identification. BMC Bioinformatics 11, 119 (2010).

54. Lander, E. S. & Waterman, M. S. Genomic mapping by fingerprinting random clones: A mathematical analysis. Genomics 2, 231–239 (1988).

