## Supplementary Material for "Performance characteristics of next-generation sequencing for antimicrobial resistance gene detection in genomes and metagenomes"

**Supplementary Figures**

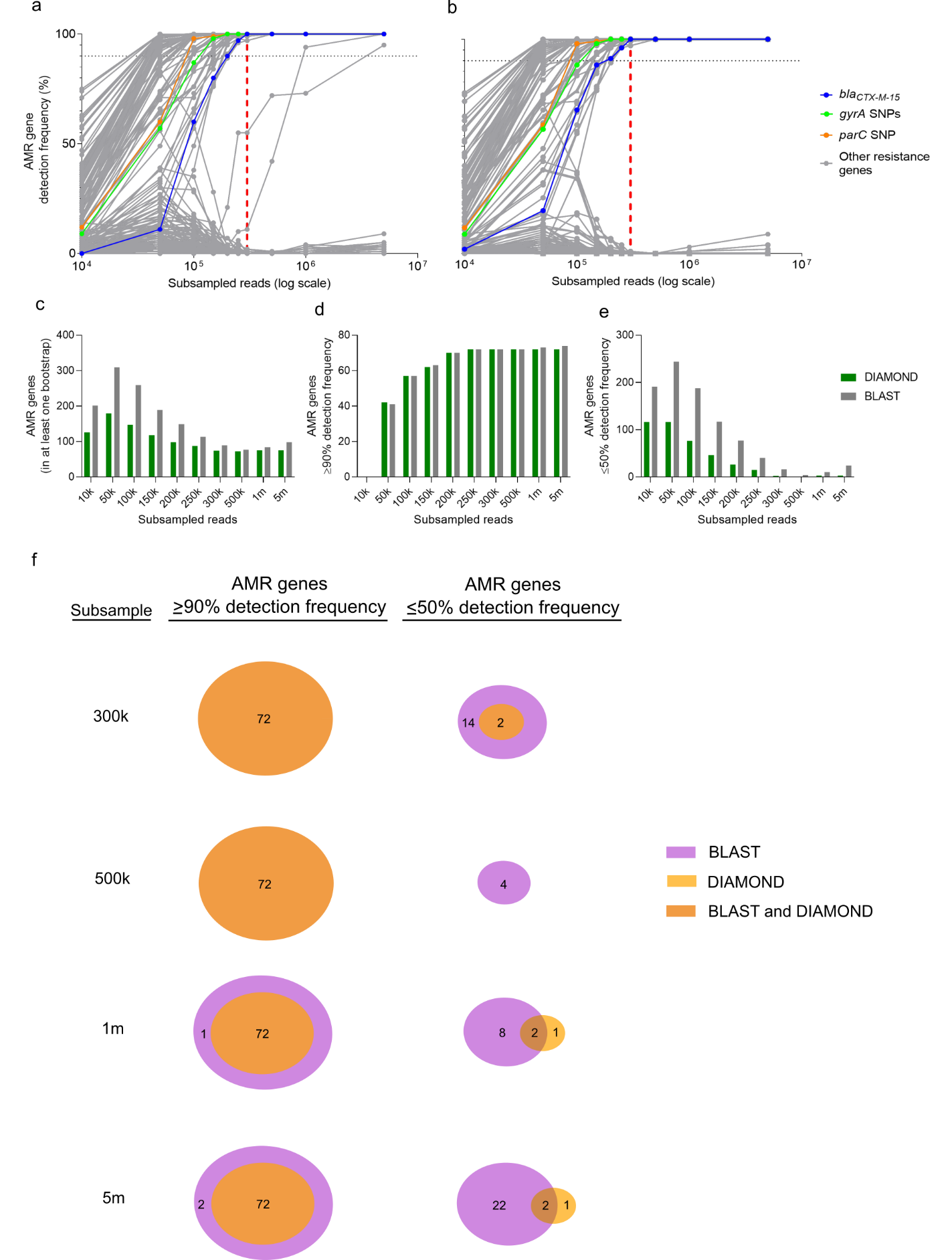

**Supplementary Figure 1.** *Escherichia coli* ST38 AMR genes detected across subsamples using either BLAST (**a**) or DIAMOND (**b**) as an aligner. Individual dots represent a single AMR gene and are connected by lines to demonstrate trends in detection across subsamples. *bla_CTX-M-15_*, *gyrA* (S83L, D87N) and *parC* (S80I) SNPs are highlighted as previously identified resistance determinants for this strain. The horizontal dotted line marks 90% detection frequency. The red vertical dashed line marks the subsample at 300,000 reads. The total number of unique genes present in 1 or more bootstraps (**c**), as well as the number of unique AMR genes predicted with ≥90% detection frequency (**d**), and with ≤50% detection frequency (**e**) using BLAST or DIAMOND as an aligner across subsamples are plotted. **f**, Venn diagrams of the number of AMR genes that were uniquely predicted by BLAST or DIAMOND or similarly predicted by BLAST and DIAMOND with ≥90% detection frequency or ≤50% detection frequency.

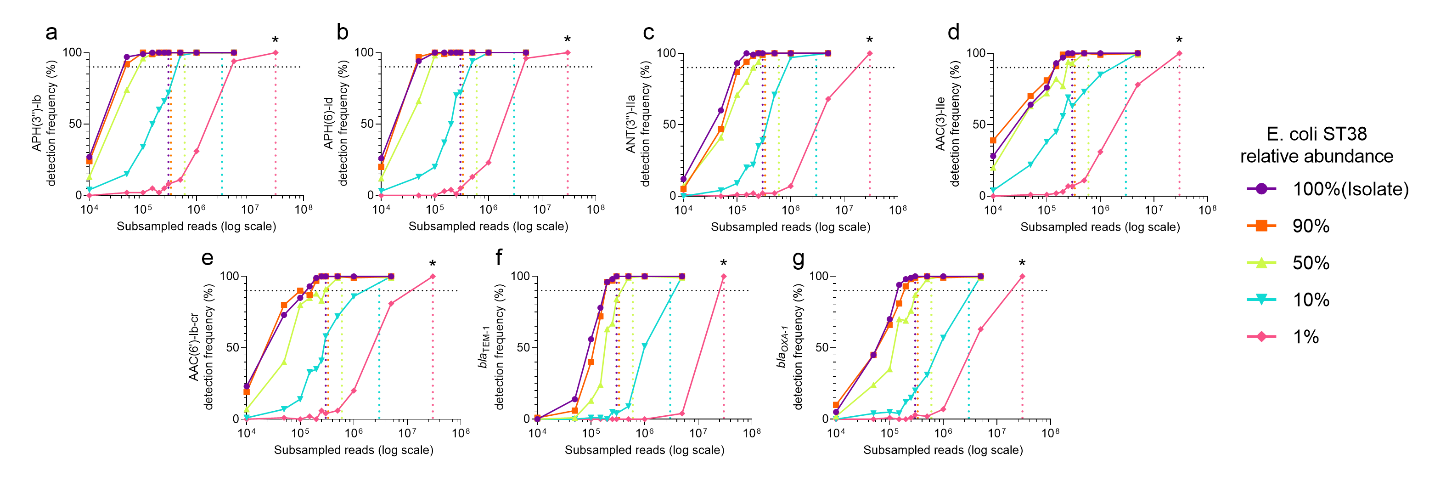

**Supplementary Figure 2.** Detection of aminoglycoside transferases (**a-e**), *bla_TEM-1_* (**f**), and *bla_OXA-1_* (**g**) from *E. coli* ST38 isolate (100%) across subsamples at varying strain relative abundances (90%, 50%, 10%, 1%) in a complex metagenomic sample using an assembly-based approach (metaSPAdes). The horizontal dotted line marks 90% detection frequency. The vertical dotted lines represent the read depths where each gene was estimated to be detected with ≥90% detection frequency for each colour matched relative abundance. The estimated read depths are 300,000, 333,333, 600,000, 3,000,000, and 30,000,000 to detect AMR genes with ≥90% detection frequency when *E. coli* ST38 is at 100%, 90%, 50%, 10%, and 1% relative abundance respectively. For all subsamples 100 bootstraps were performed, except for the 30,000,000 read subsample where 10 bootstraps were performed, which is marked by an asterisk (*). The 30,000,000 read subsample was performed on one combined sample where *E. coli* represented 1% relative abundance.

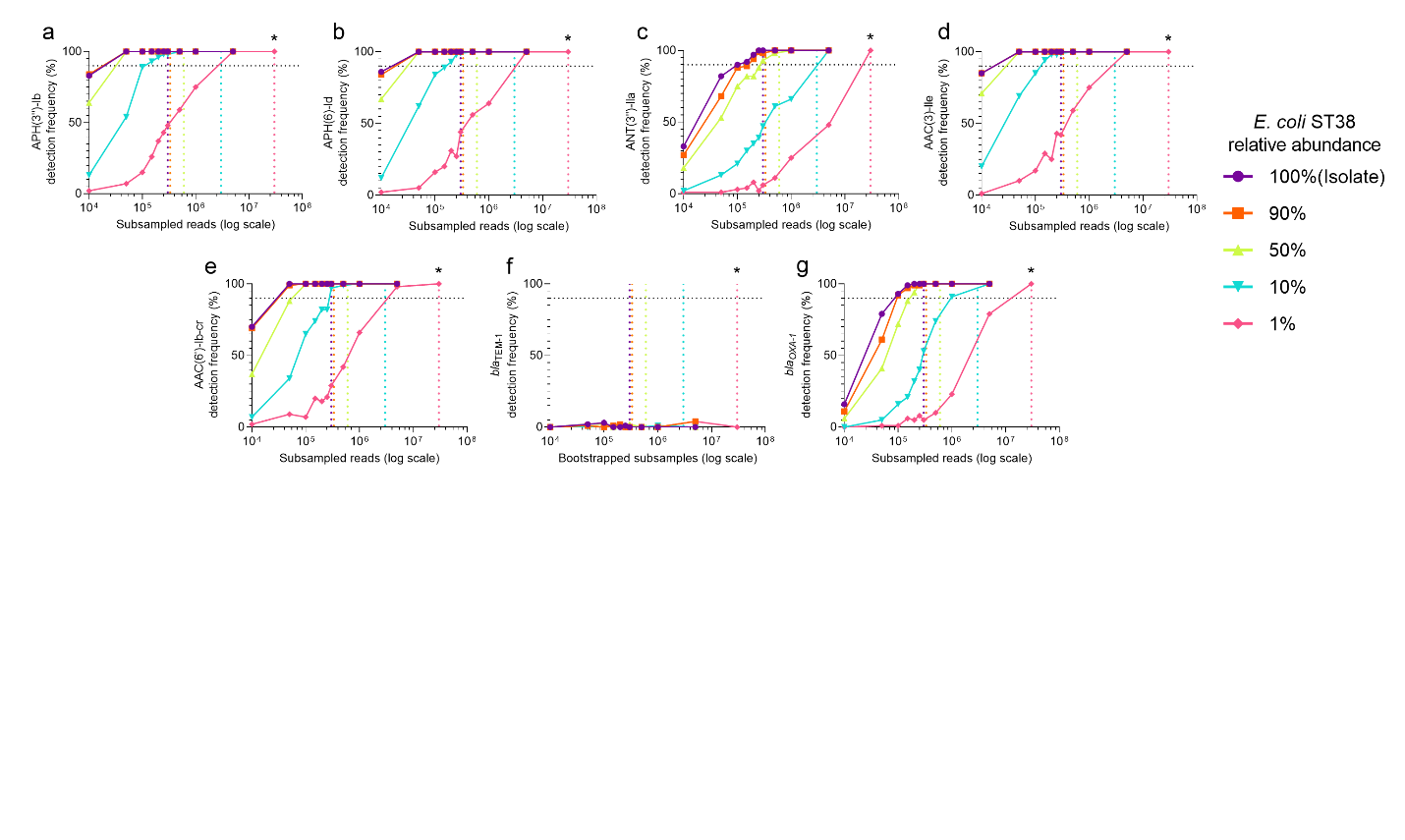

**Supplementary Figure 3.** Detection of aminoglycoside transferases (**a-e**), *bla_TEM-1_* (**f**), and *bla_OXA-1_* (**g**) from *E. coli* ST38 isolate (100%) across subsamples at varying strain relative abundances (90%, 50%, 10%, 1%) in a complex metagenomic sample using a read-based approach (KMA). The horizontal dotted line marks 90% detection frequency. The vertical dotted lines represent the read depths where each gene was estimated to be detected with ≥90% detection frequency for each colour matched relative abundance. The estimated read depths are 300,000, 333,333, 600,000, 3,000,000, and 30,000,000 to detect AMR genes with ≥90% detection frequency when *E. coli* ST38 is at 100%, 90%, 50%, 10%, and 1% relative abundance respectively. For all subsamples 100 bootstraps were performed, except for the 30,000,000 read subsample where 10 bootstraps were performed, which is marked by an asterisk (*). The 30,000,000 read subsample was performed on one combined sample where *E. coli* represented 1% relative abundance.

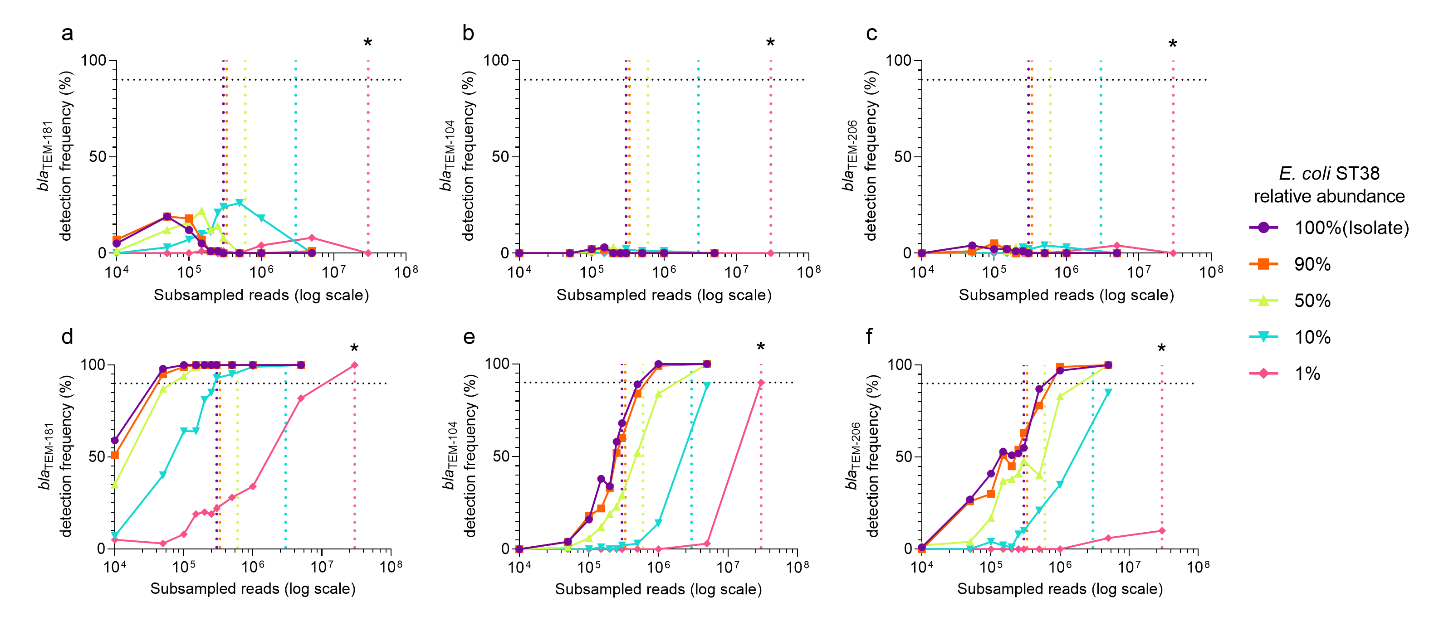

**Supplementary Figure 4.** Detection of *bla_TEM-181_* (**a,d**), *bla_TEM-104_* (**b,e**), and *bla_TEM-206_* (**c,f**) from *E. coli* ST38 isolate (100%) across subsamples at varying strain relative abundances (90%, 50%, 10%, 1%) in a complex metagenomic sample using metaSPAdes (**a-c**) or KMA (**d-f**). The horizontal dotted line marks 90% detection frequency. The vertical dotted lines represent the read depths where each gene was estimated to be detected with ≥90% detection frequency for each colour matched relative abundance. The estimated read depths are 300,000, 333,333, 600,000, 3,000,000, and 30,000,000 to detect AMR genes with ≥90% detection frequency when *E. coli* ST38 is at 100%, 90%, 50%, 10%, and 1% relative abundance respectively. For all subsamples 100 bootstraps were performed, except for the 30,000,000 read subsample where 10 bootstraps were performed, which is marked by an asterisk (*). The 30,000,000 read subsample was performed on one combined sample where *E. coli* represented 1% relative abundance.
